# Transient high glucose causes delayed wound healing by the DNMT1-mediated Ang-1/NF-κB pathway

**DOI:** 10.1101/2020.04.27.063198

**Authors:** Jingling Zhao, Shuai Yang, Lei Chen, Ronghua Yang, Yingbin Xu, Julin Xie, Xusheng Liu, Bin Shu, Shaohai Qi

## Abstract

The progression of diabetic complications does not halt despite termination of hyperglycemia, suggesting a “metabolic memory” phenomenon. However, whether metabolic memory exists in and affects the healing of diabetic wounds, as well as the underlying molecular mechanisms, remain unclear. In this study, we found that wound healing was delayed and angiogenesis was decreased in diabetic mice, despite normalization of glycemic control. Thus, we hypothesized that transient hyperglycemic spikes may be a risk factor for diabetic wound healing. We showed that transient hyperglycemia caused persistent damage to the vascular endothelium. Transient hyperglycemia directly upregulated DNMT1 expression, leading to the hypermethylation of Ang-1 and reduced Ang-1 expression, which, in turn, induced long-lasting activation of nuclear factor (NF)-κB and subsequent endothelial dysfunction. An in vivo study further showed that inhibition of DNMT1 promoted angiogenesis and accelerated diabetic wound healing by regulating the Ang-1/NF-κB signaling pathway. These results highlight the dramatic and long-lasting effects of transient hyperglycemic spikes on wound healing and suggest that DNMT1 is a novel target for diabetic vascular complications.

## Introduction

Diabetes mellitus is a chronic progressive disease that is associated with many complications, including microvasculopathy, which is characterized by alterations in vascular homeostasis due to endothelial dysfunction, leading to impaired angiogenesis and refractory wounds [1]. As a serious complication of diabetes, refractory wounds significantly impairs the quality of life for patients and leads to skin ulcers, nonhealing diabetic foot and eventual limb amputation [2–4]. Current therapies, such as growth factor treatment[5] and bioactive dressings[6, 7] have not been fully efficacious, and there is limited understanding as to which diabetic patients are susceptible to the development of chronic, non-healing wound[8] and why therapeutic strategies are effective in some patients but not in others. Therefore, the low- or non-response to therapies demonstrates the urgent need to develop new insights into the pathology of diabetic refractory wounds, leading to the identification of much needed new drug targets.

Strong epidemiological evidence from large-scale diabetes mellitus clinical trials has revealed that diabetic complications, including hypertension, neuropathy, retinopathy and nephropathy progress unimpeded even after glycemic control is pharmacologically achieved [9–11]. This harmful phenomenon is known as “metabolic memory” (MM), is supported by laboratory evidence [12–14] and is defined as permanent abnormalities in cell functions in response to initial exposure to hyperglycemia despite normalization of glycemic control [15]. Clinically, we have observed that there are no obvious improvements in delayed wound healing even when glycemic control is pharmacologically achieved in patients with diabetes. Thus, we hypothesized that transient hyperglycemic spikes exert a long-lasting effect on diabetic wound healing. However, whether the phenomenon of MM actually exists in and affects diabetic wound healing, as well as the mechanism underlying MM-induced endothelial dysfunction during wound healing, are still unclear.

The underlying molecular mechanisms of diabetic complications may include the involvement of the advanced glycation end products, the excess reactive oxygen species, and alterations in tissue-wide gene expression patterns[16–18]. However, the ability to sustain these complications in the absence of hyperglycemia invokes a role for the epigenetics in perpetuating these complications through MM. Epigenetic modifications, including DNA methylation, histone modification, and small noncoding RNAs, are essential in the complex interplay between the environment and gene expression. Epigenetic alterations create a persistent change that is stored as “memory” and passed on to the offspring, usually in response to microenvironmental stimuli [19], and these epigenetic alterations may be a key mechanism underlying MM and sustained vascular dysfunction despite achievement of glycemic control [16, 18, 20]. Recent in vitro investigations into the epigenome’s role in MM have documented specific hyperglycemia-induced DNA methylation modifications that persist in the MM state [21, 22]. Alterations of DNA methylation may become particularly important in the context of MM, as only DNA methylation is backed by strong mechanistic support for heritability of epigenetic changes[23, 24]. However, to date, the DNA methylation modifications associated with MM in diabetic refractory wounds has not been clearly studied and may reveal new targets that underlie compromised tissue regeneration and repair.

In this study, MM was examined as an important pathogenic factor in diabetic wound healing. We showed that transient hyperglycemia caused persistent damage to the vascular endothelium. Transient hyperglycemia directly upregulated DNMT1 expression, leading to the hypermethylation of Ang-1 and reduced Ang-1 expression, which, in turn, induced long-lasting activation of nuclear factor (NF)-κB and subsequent endothelial dysfunction. An in vivo study further showed that inhibition of DNMT1 promoted angiogenesis and accelerated diabetic wound healing by regulating the Ang-1/NF-κB signaling pathway. Our results implicate MM as an important mechanism in diabetic wound healing and suggest that blocking DNMT1 may have therapeutic value in treating diabetic vascular complications and delayed wound healing.

## Results

### Transient hyperglycemia induces delayed wound healing

Wound healing progressed slowly in the DM and MM groups (Fig. 1B). The wound closure rate was 30% in the control group, and it was less than 15% in the DM and MM groups at day 3. At day 7, the closure rate in the control group reached 60%, while it was only 15% and 17% in the DM and MM groups, respectively. The wounds in the control group reached complete closure by day 14, when there were still 41% and 37% unhealed wound areas in both the DM and MM groups, respectively (Fig. 1C).

**Fig. 1.**
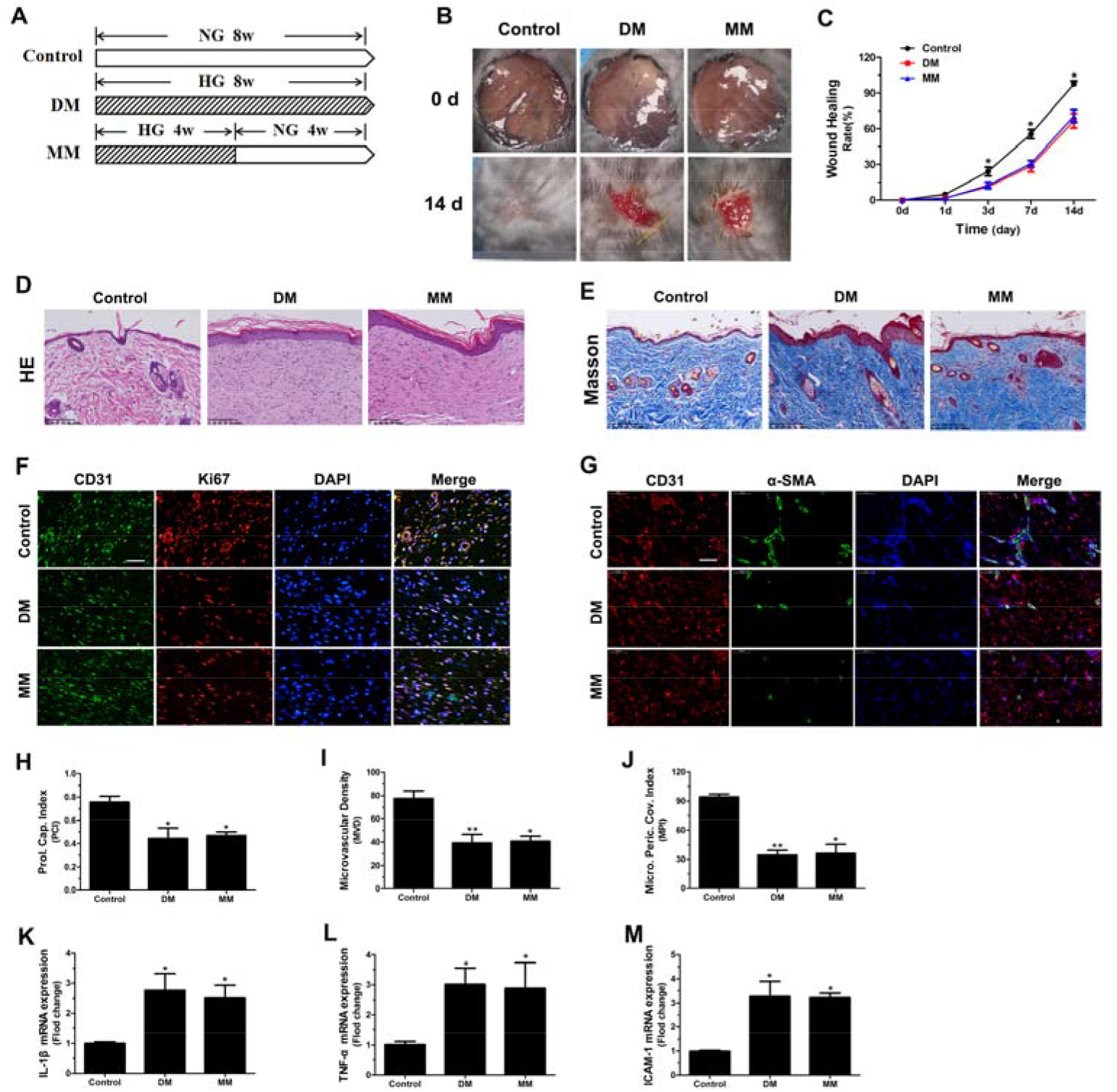
Transient hyperglycemia induces delayed wound healing. (**A**) Schematic representation of the experimental model in vivo. (**B**) Representative wound images at days 0 and 14 during the healing process. (**C**) The wound-healing rate was measured every day and is shown as the percentage of the initial wound area (n=5 per group). (**D-E**) Histological changes and collagen deposition in wounds were evaluated by H&E staining (**D**) and Masson staining (**E**). Scale bar = 100 μm. (**F-G**) Representative images of double staining of Ki67/CD31 (green, CD31; red, Ki67; nuclei, blue) (**F**) and α-SMA/CD31 (red, CD31; red, green; nuclei, blue) (**G**) in skin sections from different experimental groups. Scale bar = 50 μm. (**H-J**) Quantitative comparison of the proliferating capillary index (PCI) (**H**), microvessel density (MVD) (**I**) and microvessel pericyte coverage index (MPI) (**J**) in the different groups. (**K-M**) The expression of inflammatory cytokines, including IL-1β (**K**), TNF-α (**L**) and ICAM-1 (**M**), in the different groups. (n=10). ^*^P < 0.05, ^**^P < 0.05 (compared with control mice treated with PBS).

Histological evaluations were performed to assess the quality of wound healing. As shown in Fig. 1D, thickened epidermis, abundant infiltration of inflammatory cells and hemorrhagic scabs were observed in the DM and MM groups. Moreover, collagen deposition increased, and the collagen fibers were disorderly arranged in the DM and MM groups (Fig. 1E).

To evaluate the proliferation of endothelial cells in the wounds, PCI, which is reflected by the percentage of CD31-positive microvessels with Ki67-positive endothelial cell nuclei (Fig. 1F), was quantified, and the results showed that the PCI values were significantly lower in the DM and MM groups than in the control group (Fig. 1H). Moreover, the total number of microvessels (stained with CD31) was calculated, and the results showed that the DM and MM groups had much lower average MVDs than that of the control group (Fig. 1I). Although the MVD values reflected the presence of blood vessels, these values did not provide an indication of the maturity of the neovasculature or the functional status of the vessels. Therefore, the MPI was used to determine the percentage of microvessels covered with pericytes and was quantified to evaluate the maturity of the neovasculature (Fig. 1G). As shown in Fig. 1J, the MPIs in the DM and MM groups were observably lower than those in the control group. In addition, we detected the inflammatory condition in the wounds. As shown in Fig. 1K-M, the expression of IL-1β, TNF-α and ICAM-1, cytokines that are closely related to wound healing, markedly increased in the DM and MM groups compared with those in the control group. The above data suggest that transient hyperglycemia causes long-lasting excessive inflammatory reactions in the wound. Moreover, angiogenesis was hampered after transient hyperglycemia, which may lead to delayed wound healing and poor healing quality.

### Transient hyperglycemia damages the functions of endothelial cells

To simulate the transient hyperglycemia conditions in vitro, HUVECs were first incubated in high glucose (HG) for 24 h and then cultured in normal glucose (NG) for 2 d (MM-1), 4 d (MM-2) and 6 d (MM-3). HUVECs cultured in normal glucose were used as a control, and HUVECs cultured in high glucose alone were used to represent diabetes mellitus (DM) without tight glucose control (Fig. 2A). In addition, 30 mM mannitol served as an osmotic control (Fig S1).

**Fig. 2.**
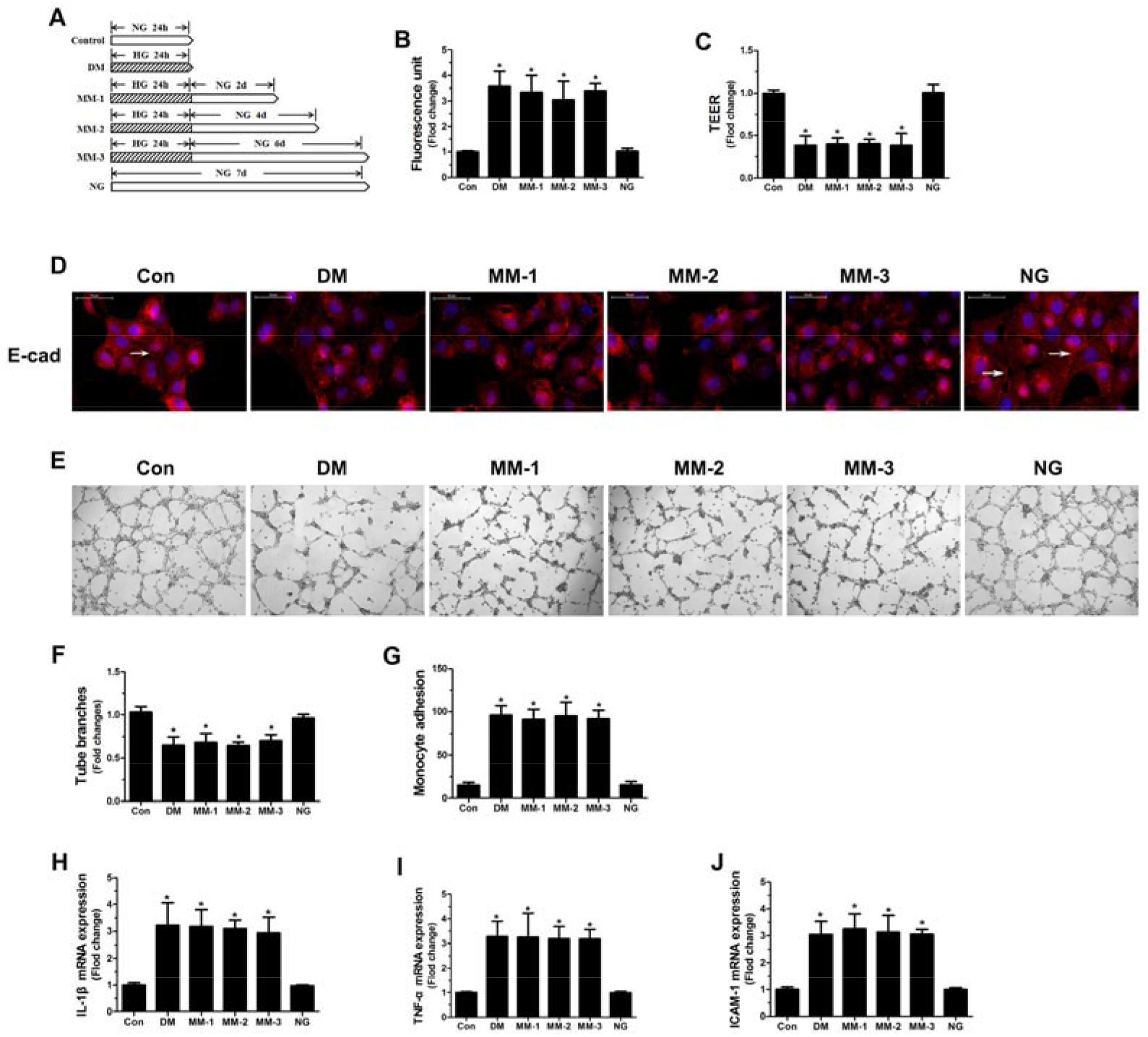
Transient hyperglycemia damages the functions of endothelial cells. (**A**) Schematic representation of the experimental model in vitro. (**B-C**) The barrier integrity of endothelial cells was evaluated using the permeability of endothelial monolayers to FITC-dextran (**B**) and the transendothelial electrical resistance (TEER) value (**C**) (n=8). (**D**) The cellular distribution of VE-cadherin (white arrows) in endothelial cells subjected to different hyperglycemia treatments. DAPI was used to stain the nuclei. Scale bar = 50 μm. (**E-F**) The tube formation assay was used to assess the ability of endothelial cells to form three-dimensional capillary-like structures in vitro and to quantify the number of tube branches (n=6). (**G**) The number of monocytes that adhered to the endothelial cells was quantified by the monocyte adhesion assay (n=6). (**H-J**) Relative mRNA expression of the inflammatory cytokines IL-1β (**H**), TNF-α (**I**) and ICAM-1 (**J**) in endothelial cells subjected to different hyperglycemia treatments (n=6). ^*^P < 0.05 (compared with control group).

We first assessed the barrier integrity of endothelial cells. As shown in Fig. 2B, HG treatment significantly increased the permeability of endothelial monolayers to FITC-dextran, which persisted in a normoglycemic environment for the entire 6-d experimental period. Correspondingly, the TEER of endothelial monolayers decreased after HG treatment and remained at this low level after subsequent exposure to NG (Fig. 2C). Given that modifications of cellular junction protein can result in endothelial monolayer hyperpermeability [25], we thus detected the presence of the junctional protein E-cadherin. We found that E-cadherin was evenly distributed in a continuous pattern along the endothelial intercellular junctions in the control group, and HG treatment disrupted cell-cell contacts and caused a substantial loss of cell surface-associated E-cadherin expression. Notably, this decrease in E-cadherin expression and interrupted expression pattern persisted during subsequent incubation under NG conditions (Fig. 2D).

Next, the tube formation ability of endothelial cells was determined. As shown in Fig. 2E, the ability of endothelial cells to form capillary-like tubes was dramatically reduced after HG treatment, and this reduction persisted for 6 d with subsequent exposure to NG. The quantification of total branching points showed that the number of tube branches significantly decreased with transient exposure to HG, which also persisted for 6 d (Fig. 2F).

Since increased adhesion molecules on endothelial cells lead to the infiltration of inflammatory leukocytes into the vessel wall, which is a hallmark of a dysfunctional endothelium [26], we investigated the effect of HG on the adhesion of monocytes to endothelial cells. The results showed that the adhesion of monocytes to endothelial cells was stimulated by transient hyperglycemia and remained elevated during 6 d of subsequent incubation under NG conditions (Fig. 2G).

Finally, the expression of the proinflammatory cytokines IL-1β, TNF-α and ICAM-1 was detected. Similar to that observed with monocyte adhesion, transient hyperglycemia induced an increase in the expression of these cytokines, which persisted for the subsequent 6 d of exposure to NG conditions (Fig. 2H-J).

The above data suggest that transient hyperglycemia causes persistent injuries to the functions of endothelial cells, including barrier integrity and tube formation ability, and long-lasting excessive inflammation may be involved in the damage process.

### Transient hyperglycemia damages the functions of endothelial cells by inhibiting the expression of Ang-1

Our previous research reported that Ang-1 plays an important role in advanced glycation end product (AGE)-induced endothelial cell dysfunction [27]. AGEs are modifications of proteins or lipids that are prevalent in the diabetic vasculature and are considered to be an important cause of diabetic vascular complications [28]; therefore, we asked whether Ang-1 is implicated in transient hyperglycemia-induced persistent microvascular impairments. As shown in Fig. 3A-C, the mRNA and protein expression of Ang-1 significantly decreased after exposure to transient HG, and this decrease persisted for 6 d of subsequent exposure to NG conditions.

**Fig. 3.**
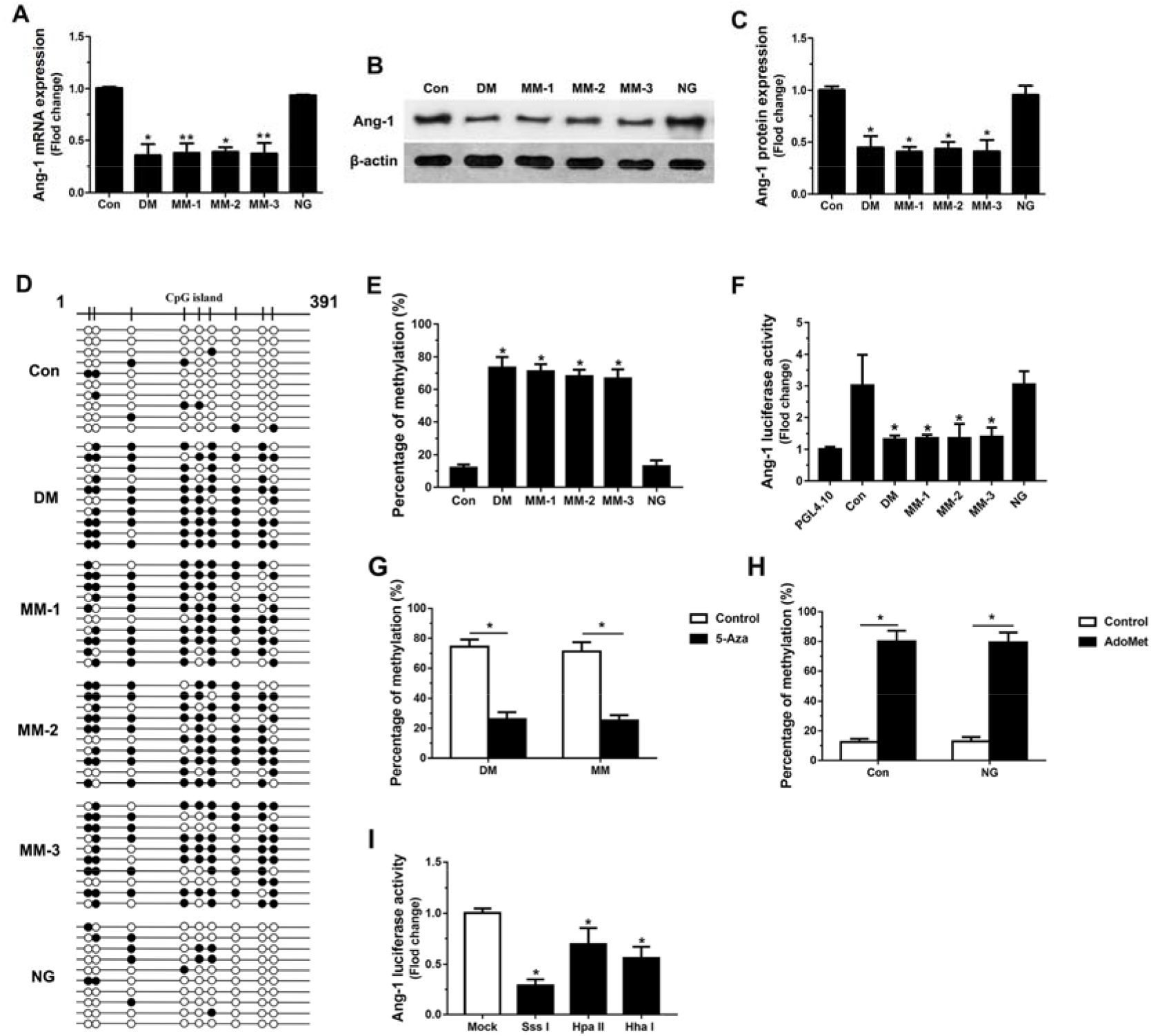
The transient hyperglycemia-induced decrease in Ang-1 expression is linked to hypermethylation of the Ang-1 promoter. (**A**) The mRNA expression of Ang-1 in endothelial cells subjected to different hyperglycemia treatments (n=6). (**B-C**) Quantification of Ang-1 expression by western blot analysis (n=4). (**D-E**) BSP analysis of the Ang-1 promoter in endothelial cells subjected to different hyperglycemia treatments (n=4). (**F**) Ang-1 promoter activity in endothelial cells (n=6). (**G-H**) BSP analysis of the Ang-1 promoter in endothelial cells after treatment with 5-Aza or DAC (n=4). (**I**) Ang-1 promoter activity in endothelial cells after Sss I, Hpa II and Hha I treatment (n=6). ^*^P < 0.05 (compared with control group).

To further investigate the effect of Ang-1 on endothelial cells subjected to transient hyperglycemia injury, a potent recombinant Ang-1 protein variant, cartilage oligomeric matrix protein (COMP)-angiopoietin-1 (cAng-1), was used [29]. The results showed that cAng-1 treatment revered the transient HG-induced high permeability and low TEER of endothelial monolayers (Fig. S2A-B). Similarly, cAng-1 treatment prevented the substantial loss of cell surface-associated E-cadherin expression that was caused by transient hyperglycemia (Fig. S2C). Moreover, cAng-1 improved the transient HG-mediated injury to the tube formation ability of endothelial cells and abrogated the reduction in the number of tube branches (Fig. S2D-E). In addition, cAng-1 observably inhibited the transient HG-induced persistent excessive inflammatory conditions, including the increased adhesion of monocytes to endothelial cells (Fig. S2F) and overexpression of IL-1β, TNF-α and ICAM-1 (Fig. 2G-I). These data indicate that transient hyperglycemia damages the functions of endothelial cells by inhibiting the expression of Ang-1.

### Decreased Ang-1 expression induced by transient hyperglycemia is linked to hypermethylation of the Ang-1 promoter

We next studied how transient hyperglycemia affects the expression of Ang-1. As shown in Fig. 3D-E, when compared with the methylation levels in endothelial cells cultured under normal conditions, increased methylation was detected by bisulfite sequencing PCR (BSP) in the promoter region of Ang-1 in cells after transient hyperglycemia, and this increase persisted for 6 d of subsequent exposure to NG conditions. Moreover, Ang-1 promoter activity was significantly decreased in transient hyperglycemia-treated endothelial cells, which also persisted for 6 d of subsequent exposure to NG conditions (Fig. 3F). Furthermore, treatment of endothelial cells with the methyltransferase inhibitor 5-aza-deoxycytidine (5-Aza), an effective DNA demethylating agent, significantly decreased the methylation level of the Ang-1 promoter and resulted in activation of Ang-1 gene expression (Fig. 3G). Conversely, treatment of cells with the methyl donor S-adenosyl-L-methionine (AdoMet), which has been shown to inhibit demethylase activity and induce DNA methylation [30], led to upregulated methylation of the Ang-1 promoter (Fig. 3H). To further explore whether DNA methylation directly regulates Ang-1 promoter activity, the promoter region of Ang-1 was cloned into a luciferase reporter construct. SssI methylase (methylation of 12 CpGs), HpaII methylase (methylation of 2 CpGs), and HhaI methylase (methylation of 2 CpGs) [31] were used to clone the fragment, and Ang-1 promoter activity was determined after transfection of endothelial cells with methylated or mock-methylated luciferase constructs. As shown in Fig. 3I, methylation inhibited Ang-1 promoter activity in a methylation dose–dependent manner. Taken together, the above results suggest that decreased Ang-1 expression is associated with increased methylation of the Ang-1 promoter.

### Elevated DNMT1 expression is involved in the transient hyperglycemia-induced decrease in Ang-1 expression

We demonstrated that aberrant DNA methylation is associated with decreased Ang-1 expression. DNA methylation is catalyzed by DNA methyltransferases (DNMTs): DNMT3a and 3b are de novo enzymes, and DNMT1 is a maintenance enzyme that is important in regulating tissue-specific patterns of methylated cytosine [32]. Here, we detected the expression of DNMTs, and the results showed that the mRNA and protein expression of DNMT1 were increased (Fig. 4A-C) and the DNMT1 activity was elevated (Fig. 4D) in endothelial cells that were subjected to HG injury, and the increased status persisted during subsequent incubation under NG conditions. Immunofluorescence staining also showed that DNMT1 was mainly located in the nucleus, and the expression remained high after transient HG treatment (Fig. 4E). However, neither the activity nor expression of DNMT3a nor 3b changed in endothelial cells after HG treatment (Fig. S3). In addition, the ChIP assay indicated that the binding of DNMT1 to the Ang-1 promoter was enhanced by transient HG treatment, and these changes persisted for 6 d of subsequent incubation at normal glucose levels (Fig. 4F).

**Fig. 4.**
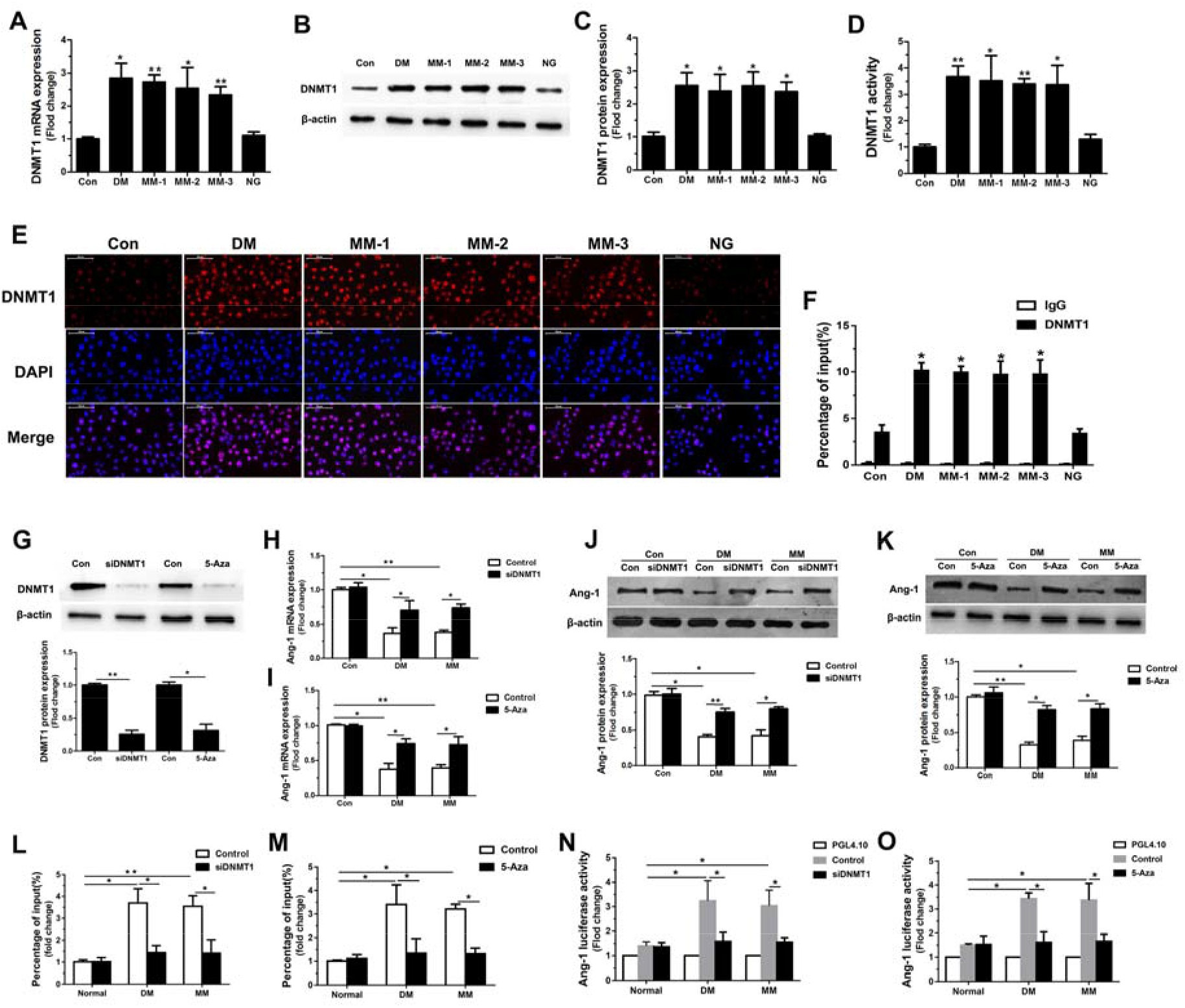
Elevated DNMT1 expression is involved in the transient hyperglycemia-induced decrease in Ang-1 expression. (**A**) The mRNA expression of DNMT1 in endothelial cells subjected to different hyperglycemia treatments (n=6). Quantification of the protein expression of DNMT1 (n=4) (**B-C**) and the activity of DNMT1 (n=6) (**D**). (**E**) Immunofluorescence staining showing the expression and cellular localization of DNMT1 in endothelial cells subjected to different hyperglycemia treatments. Scale bar = 50 μm. (**F**) ChIP assay showing the binding of DNMT1 to the promoter of Ang-1 in endothelial cells (n=6). (**G**) DNMT1 expression was knocked down by siRNA and 5-Aza in endothelial cells, and the efficiency of knockdown was confirmed by western blot analysis (n=4). (**H-I**) Relative mRNA expression of Ang-1 in endothelial cells that with DNMT1 knockdown under different hyperglycemia conditions (n=6). (**J-K**) Western blot analysis showing the expression Ang-1 in endothelial cells with DNMT1 knockdown by siDNMT1 (**J**) or 5-Aza (**K**) (n=4). (**L-M**) The binding of DNMT1 to the promoter of Ang-1 in endothelial cells after inhibition of DNMT1 (n=6). (**N-O**) Luciferase reporter assay showing the Ang-1 promoter activity of endothelial cells after inhibition of DNMT1 (n=6). ^*^P < 0.05 (compared with control group).

To clarify whether DNMT1 directly regulated Ang-1 expression, the specific DNMT1 inhibitor 5-Aza and siRNA against DNMT1 (siDNMT1) were used to inhibit the expression of DNMT1, and the efficiency of inhibition was confirmed by western blot analysis (Fig. 4G). The results showed that siDNMT1 or 5-Aza treatment reversed the decreased mRNA (Fig. 4H-I) and protein expression of Ang-1 that was induced by transient HG exposure (Fig. 4J-K). Moreover, after inhibition of DNMT1 expression, the binding of DNMT1 to the Ang-1 promoter was decreased (Fig. 4L-M), while Ang-1 promoter activity was enhanced even after transient HG treatment (Fig. 4N-O). These data suggest that transient HG-induced upregulation of DNMT1 directly represses Ang-1 activation and expression.

### Elevated DNMT1 expression is involved in transient hyperglycemia-induced endothelial cell dysfunction

We demonstrated that transient hyperglycemia damages the functions of endothelial cells by inhibiting the expression of Ang-1, while DNMT1 directly regulates Ang-1 activation and expression. Thus, we asked whether DNMT1 is involved in transient hyperglycemia-induced endothelial cell dysfunction. As shown in Fig. 5A-D, siDNMT1 or 5-Aza treatment reversed the transient HG-induced high permeability and low TEER value of endothelial monolayers. Moreover, the addition of 5-Aza or siDNMT1 application largely restored the loss of cell surface-associated E-cadherin expression caused by transient hyperglycemia treatment (Fig. 5E-F). Also, siDNMT1 or 5-Aza treatment improved the transient HG-induced weakened tube formation ability of endothelial cells (Fig. 5G-J). In addition, siDNMT1 or 5-Aza treatment significantly suppressed the excessive inflammatory response, including the increased adhesion of monocytes to endothelial cells (Fig. 5K-L) and overexpression of IL-1β, TNF-α and ICAM-1, caused by transient hyperglycemia treatment (Fig. 5M-R). These data indicate that during transient hyperglycemia, DNMT1 induced long-lasting damage to endothelial cells by inhibiting Ang-1 activation and expression.

**Fig. 5.**
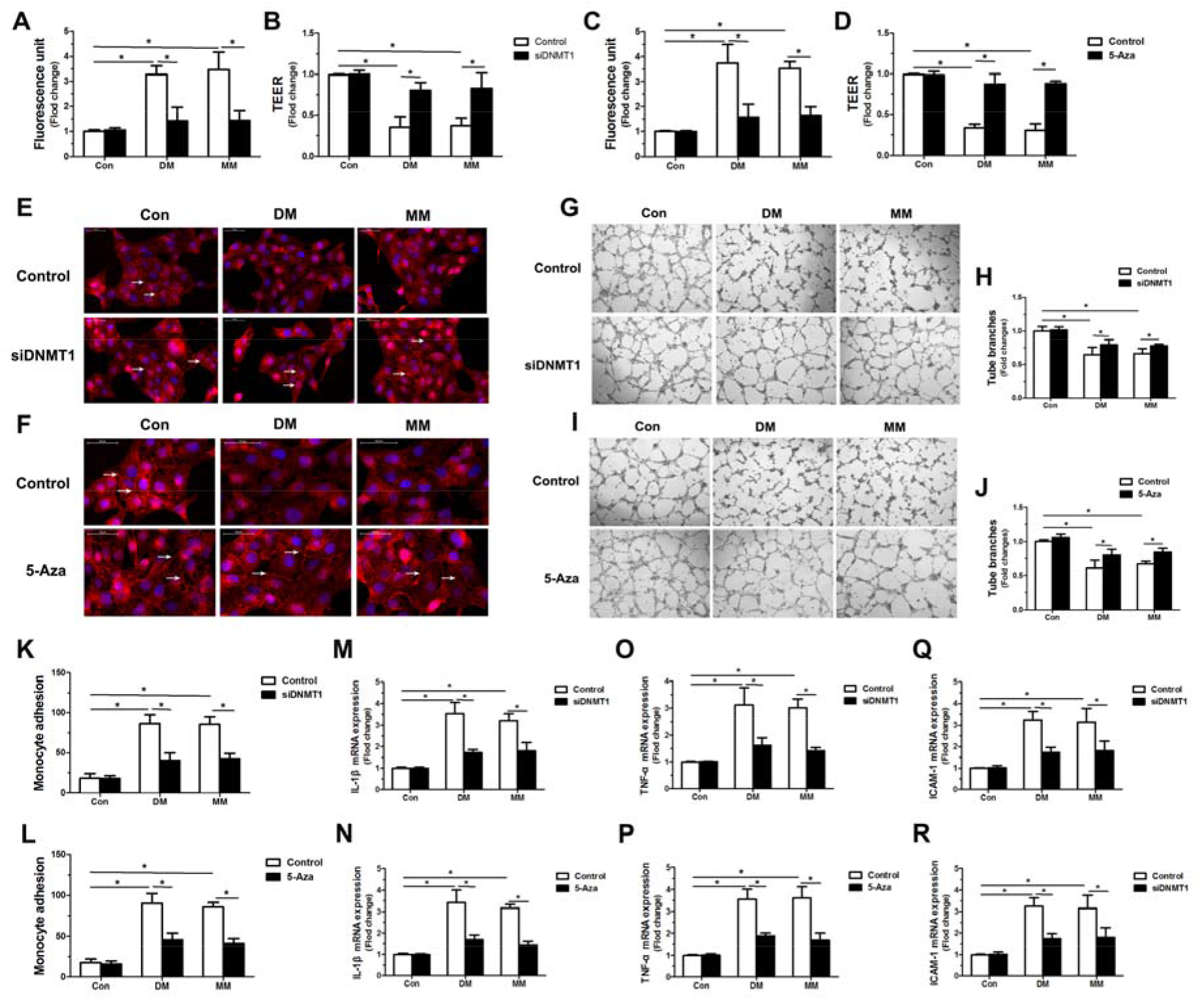
Elevated DNMT1 expression is involved in transient hyperglycemia-induced endothelial cell dysfunction. (**A-D)** The permeability of endothelial monolayers to FITC-dextran and the TEER value of endothelial cells treated with siDNMT1 (**A-B**) or 5-Aza (**C-D**) under different hyperglycemia conditions (n=8). (**E-F**) The cellular distribution of VE-cadherin in endothelial cells after inhibition of DNMT1 with siDNMT1 (**E**) or 5-Aza (**F**). Scale bar = 50 μm. (**G-J**) The tube formation ability of endothelial cells treated with siDNMT1 (**G-H**) or 5-Aza (**I-J**) (n=8). (**K-L**) The number of monocytes that adhered to the endothelial cells after inhibition of DNMT1 with siDNMT1 (**K**) or 5-Aza (**L**). (n=6). (**M-R**) Relative mRNA expression of the inflammatory cytokines IL-1β (**M-N**), TNF-α (**O-P**) and ICAM-1 (**Q-R**) in endothelial cells treated with siDNMT1 or 5-Aza (n=6). ^*^P < 0.05 (compared with control group).

### NF-κB pathway activation is involved in DNMT1/Ang-1 signaling-induced endothelial cell dysfunction

We found that DNMT1/Ang-1 signaling is required for transient hyperglycemia-induced endothelial cell dysfunction. We next explored the underlying mechanisms by which DNMT1/Ang-1 signaling influences endothelial cell function. Ang-1 functions as a strong endothelial-specific protective factor and protects endothelial cell function by inhibiting the activation of the NF-κB pathway [33, 34]. Active methylation and increased NF-κB transactivation play critical roles in the development of diabetic complications, such as nephropathy and retinopathy [35, 36], suggesting that the effect of DNMT1/Ang-1 signaling on transient hyperglycemia-induced endothelial cell dysfunction involves the NF-κB pathway. Therefore, we aimed to determine whether DNMT1/Ang-1 affects endothelial cell function by regulating the NF-κB pathway.

We initially examined the effect of different hyperglycemia conditions on NF-κB signaling. The results showed that the phosphorylation of NF-κB p65 (at Ser536) increased after transient hyperglycemia treatment, and p-p65 expression remained at this high level after subsequent exposure to normal glucose (Fig. 6A). Immunofluorescence staining showed that there was a significant increase in the nuclear translocation of the NF-κB subunit p65 in cells after transient hyperglycemia stimulation, and this nuclear distribution pattern persisted during subsequent incubation under normal glucose conditions (Fig. 6B-C). Moreover, a luciferase reporter assay also showed that transient hyperglycemia induced an increase in NF-κB luciferase reporter activity, which persisted for the subsequent 6 d of exposure to normal glucose conditions (Fig. 6D).

**Fig. 6.**
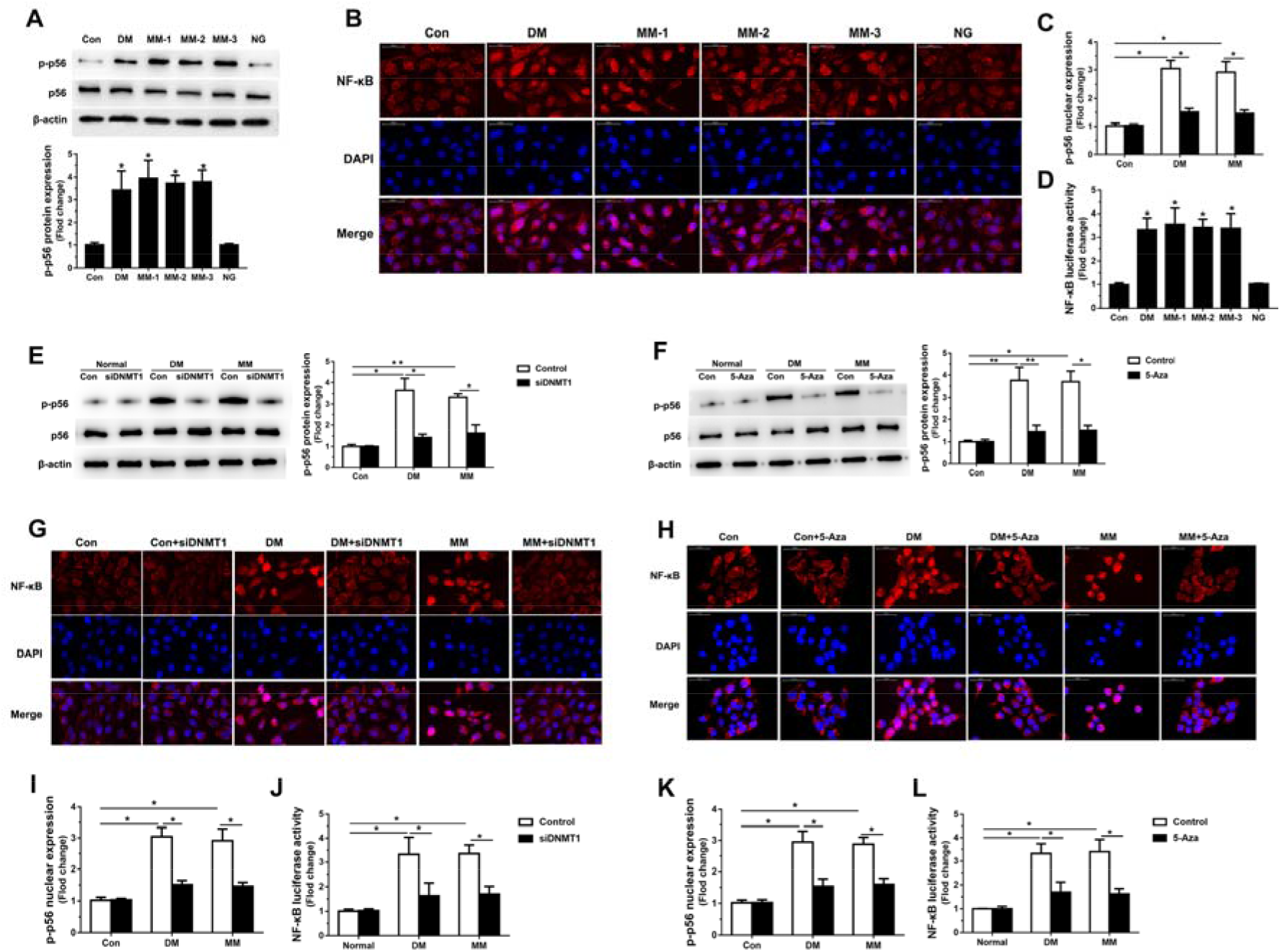
Transient hyperglycemia activates the NF-κB pathway by increasing DNMT1 expression. (**A**) Western blot analysis showing the phosphorylation of NF-κB p65 (at Ser536) in endothelial cells subjected to different hyperglycemia treatments (n=4). (**B-C**) Immunofluorescence staining showing the expression and cellular localization of NF-κB in endothelial cells. Scale bar = 50 μm. (**D**) Luciferase reporter activity of NF-κB in endothelial cells subjected to different hyperglycemia treatments (n=6). (**E-F**) The phosphorylation of NF-κB p65 in endothelial cells after inhibition of DNMT1 with siDNMT1 (**E**) or 5-Aza (**F**) (n=4). (**G-H**) The expression and cellular localization of NF-κB in endothelial cells after inhibition of DNMT1 with siDNMT1 (**G**) or 5-Aza (**H**). Scale bar = 50 μm. (**I, K**) Quantification of the nuclear expression of NF-κB. (**J, L**) Luciferase reporter activity of NF-κB in endothelial cells after inhibition of DNMT1 with siDNMT1 (**J**) or 5-Aza (**L**) (n=6). ^*^P < 0.05 (compared with control group).

We next investigated whether DNMT1 directly affects NF-κB activity. As shown in Fig. 6E-F, inhibition of DNMT1 expression with siDNMT1 or 5-Aza significantly decreased the transient hyperglycemia-induced phosphorylation of NF-κB p65. In addition, cells with reduced DNMT1 expression showed reduced nuclear NF-κB p65 expression (Fig. 6G-H, I and K) and reduced NF-κB luciferase reporter activity (Fig. 6J, L), even in hyperglycemia conditions.

We then examined whether Ang-1 was essential for transient hyperglycemia-induced NF-κB activation. The results showed that transient hyperglycemia-induced phosphorylation of NF-κB p65 (Fig. S4A), nuclear translocation of NF-κB p65 (Fig. S4B-C), and increased NF-κB luciferase reporter activity (Fig. S4D) were markedly reversed by cAng-1 treatment.

Finally, we determined whether blocking the NF-κB pathway using the NF-κB inhibitor BAY 11-7085, which prevents the nuclear translocation of p65/p50 [37], attenuated transient hyperglycemia-induced endothelial cell dysfunction. As shown in Fig, S5A-B, BAY 11-7085 treatment reversed the transient hyperglycemia-induced high permeability and low TEER value of endothelial monolayers. BAY 11-7085 also prevented the loss of cell surface-associated E-cadherin expression that was caused by transient hyperglycemia (Fig. S5C). Moreover, BAY 11-7085 treatment observably inhibited the transient HG-injured tube formation ability of endothelial cells (Fig. S5D-E), as well as the excessive inflammatory responses (Fig. S5F-I).

Taken together, our findings indicate that DNMT1/Ang-1 affects endothelial cell functions during transient hyperglycemia by regulating the NF-κB pathway.

### Inhibition of DNMT1 promotes the healing of diabetic wounds

To determine in vivo whether DNMT1 and its downstream effects play important roles in hyperglycemia (DM) or transient hyperglycemia (MM) induction of delayed wound healing, we initially examined DNMT1 expression during wound healing. Immunohistochemical staining showed strong positive expression of DNMT1 in the wounds from the DM and MM groups. In contrast, weak staining for DNMT1 was observed in the wounds from the control group (Fig. S6A). Quantitative detection using western blotting showed that when compared with the expression level in the control group, the expression level of DNMT1 was significantly increased in the DM and MM groups (Fig. S6B). Furthermore, the expression of Ang-1 was clearly reduced (Fig. S6C-D), and the activation of NF-κB was increased in the DM and MM groups when compared with those in the control group (Fig. S6E). These findings suggest that the aberrant expression of DNMT1 and its downstream effects are closely related to metabolic memory-induced delayed wound healing.

To further clarify whether inhibition of DNMT1 can rescue the delayed wound healing caused by DM or MM, siDNMT1 or 5-Aza diluted in PBS was applied to the wounds, and PBS was used as a control. We found that the wound closure rate was significantly increased in the DM and MM groups that were treated with either siDNMT1 or 5-Aza (Fig. 7A-B). Histological analysis also showed that inhibition of DNMT1 by siDNMT1 or 5-Aza improved the reepithelialization and revascularization, as well as decreased the excessive inflammatory cell infiltration and collagen deposition during wound healing (Fig. 7C). Moreover, the repressed angiogenic process in the DM and MM groups (Fig. 7D-E), such as decreased proliferation of endothelial cells (PCI) (Fig. 7G), reduced number of microvessels (MVD) (Fig. 7F) and the immaturity of the neovasculature (MPI) (Fig. 7H), were significantly improved after inhibition of DNMT1 by siDNMT1 or 5-Aza. The overexpression of inflammatory cytokines, including IL-1β, TNF-α and ICAM-1, in the DM and MM groups was obviously reduced by siDNMT1 or 5-Aza treatment. (Fig. 7I-K). In addition, the increased expression of DNMT1 in the DM and MM groups was reduced after siDNMT1 or 5-Aza treatment (Fig. 8A-B). More importantly, the decreased expression of Ang-1 (Fig. 8C-D) and the upregulated phosphorylation of NF-κB p65 (Fig. 8E) were reversed after inhibition of DNMT1 by siDNMT1 or 5-Aza.

**Fig. 7.**
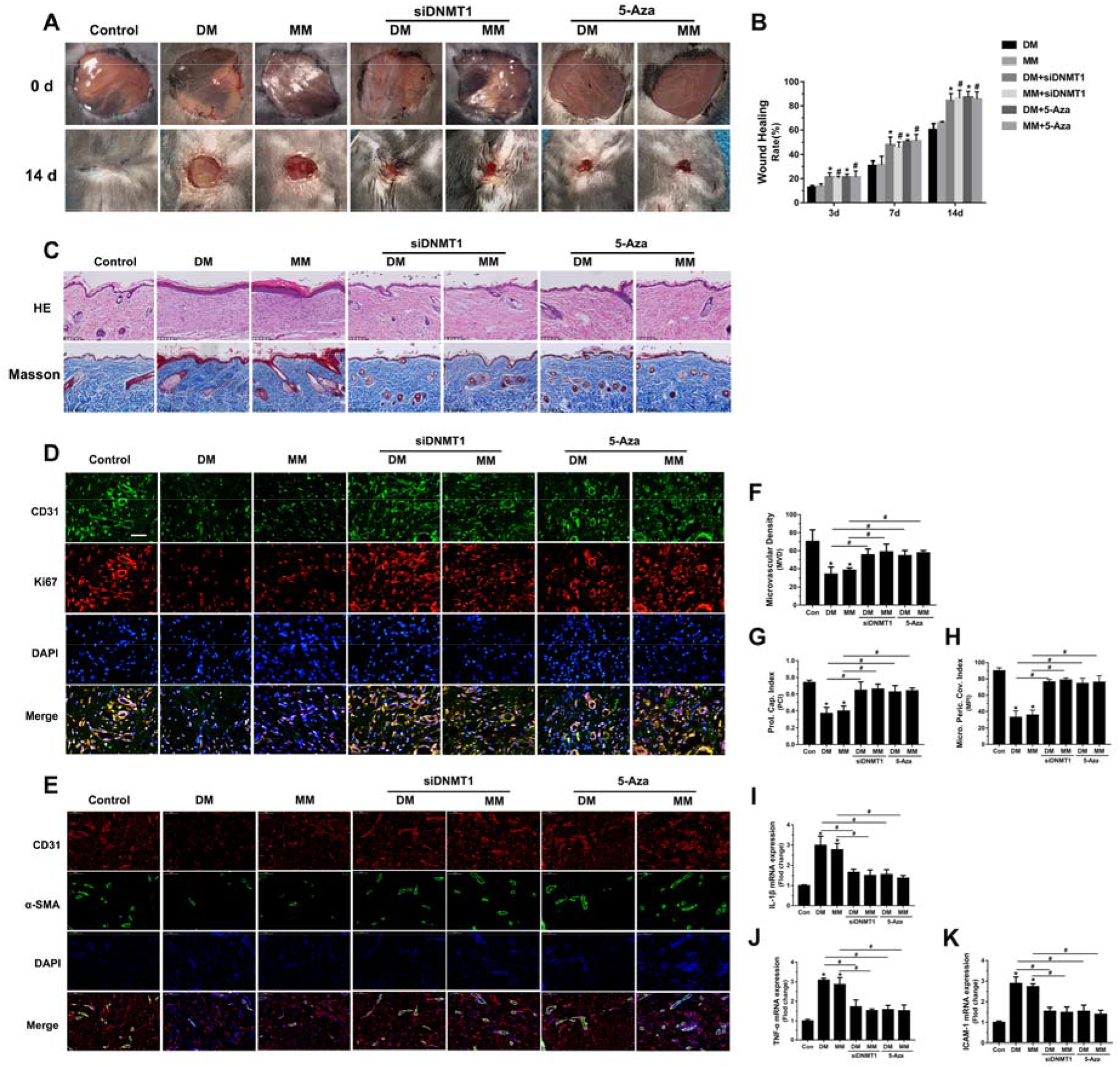
Inhibition of DNMT1 promotes the healing of diabetic wounds. (**A**) Representative wound images at days 0 and 14 during the healing process. (**B**) The wound-healing rate was quantified and is shown as the percentage of the initial wound area (n=5 per group). (**C**) Histological changes and collagen deposition in wounds were evaluated by H&E and Masson staining. Scale bar = 100 μm. (**D-E**) Representative images of double staining of Ki67/CD31 (**D**) and α-SMA/CD31 (**E**) in skin sections. Scale bar = 50 μm. (**C-E**) Quantitative comparison of the proliferating capillary index (PCI) (**G**), microvessel density (MVD) (**F**) and microvessel pericyte coverage index (MPI) (**H**) in the different groups. (**I-K**) The expression of inflammatory cytokines, including IL-1β (**I**), TNF-α (**J**) and ICAM-1 (**K**), in the different groups. (n=10). ^*^P < 0.05 (compared with control mice treated with PBS), ^#^P < 0.05.

**Fig. 8.**
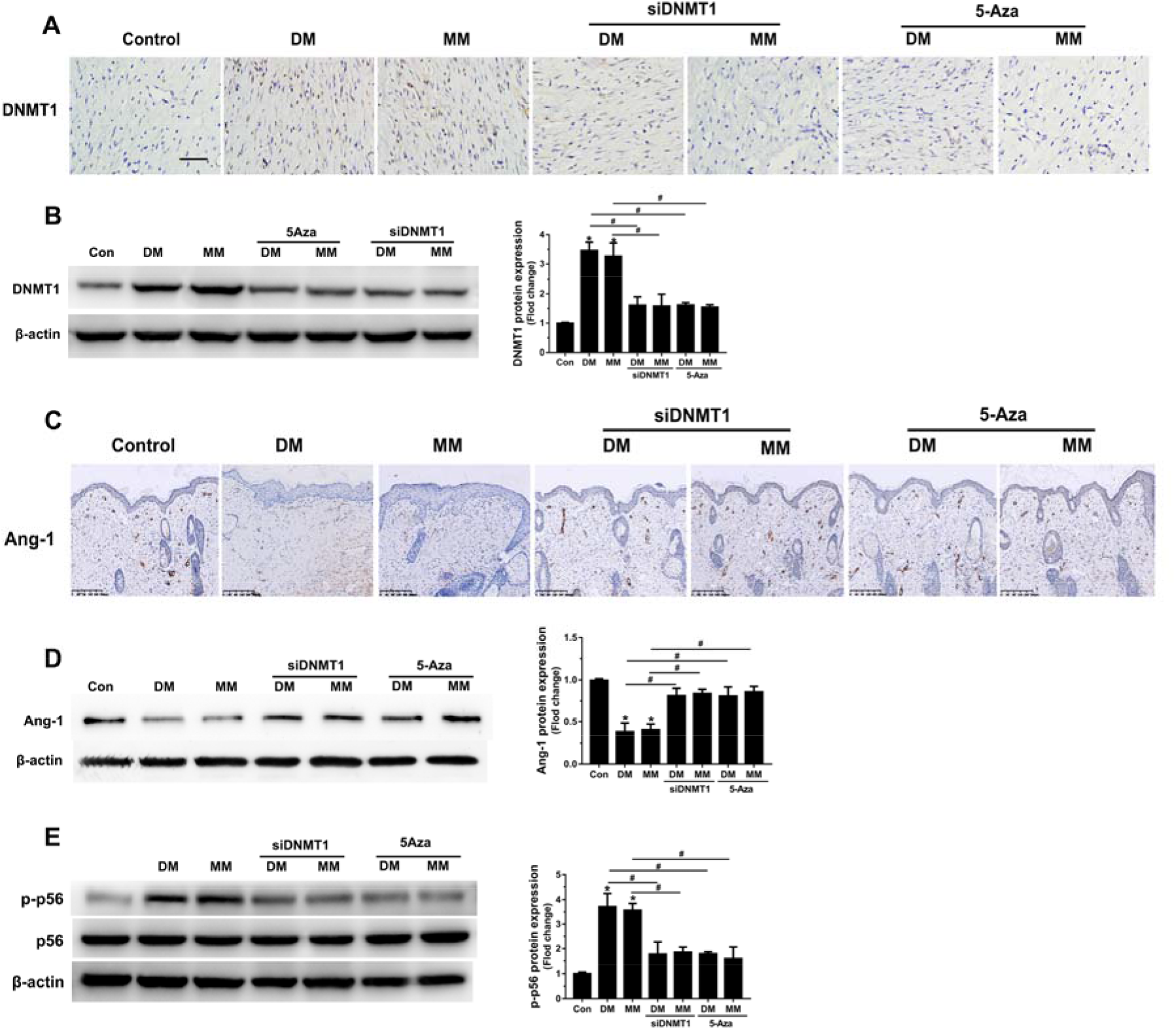
Inhibition of DNMT1 affects the expression of Ang-1/NF-κB during the healing of diabetic wounds. (**A**) Immunohistochemical staining showing the expression of DNMT1 in the wounds that were subjected to different treatments. Scale bar = 100 μm. (**B**) Western blot analysis showing the expression of DNMT1 in the wounds (n=4). (**C**) Immunohistochemical staining showing the expression of Ang-1 in the wounds. Scale bar = 200 μm. (**D**) Western blot analysis showing the expression of Ang-1 in the wounds (n=4). (**E**) Western blot analysis showing the phosphorylation of NF-κB p65 in the wounds (n=4). ^*^P < 0.05 (compared with control mice treated with PBS), ^#^P < 0.05.

These data indicate that inhibition of DNMT1 can accelerate diabetic wound healing by promoting vascularization and inhibiting inflammatory responses, and the Ang-1/NF-κB pathway serves as the potential mechanism.

## Discussion

Our findings are particularly novel for two reasons. First, to our knowledge there are no detailed research about whether metabolic memory plays an important role in the diabetic wound healing. Here we found that wound healing progressed slowly in diabetic mice despite achievement of glycemic control, indicating the “metabolic memory” phenomenon indeed exists in and affects the healing of diabetic wounds. The persistent impairment in angiogenesis induced by transient hyperglycemia exposure is a major reason for why diabetic patients are susceptible to the development of chronic, non-healing wound and why conventional therapeutic strategies are not effective in these patients. Second, and most important clinically, our study is the first to demonstrate that the DNMT1-mediated aberrant Ang-1 DNA methylation and vascular epigenetic changes that caused by transient hyperglycemia is the key mechanism that result in delayed wound healing during subsequent normoglycemia.

Impaired wound healing is a serious diabetic complication with high morbidity, a long course and limited effective treatment [1]. Clinically, we have observed that many diabetic patients who already have good glycemic control continue to experience refractory wounds. In this study, we found that long-lasting excessive inflammatory reaction and deficiency in angiogenesis is due to continuing injury from exposure to transient hyperglycemic conditions, leading to delayed wound healing through metabolic memory. A central mechanism underlying metabolic memory is the hyperglycemia-induced epigenetic patterns that drive and sustain the progression of disease phenotypes[38]. Initial researches have revealed that histone modifications are maintained in the posthyperglycemic environment[39, 40]. However, DNA methylation, which has the strongest experimental support for heritability and could contribute to the heritable transmission[23], has been recently associated with metabolic memory and progression of diabetic complications. Using a diabetic zebrafish model, it was found that the reduced fin regeneration caused by hyperglycemia are maintained through the metabolic memory, and the global DNA hypomethylation that correlated with aberrant gene expression is the potential mechanism[41]. Another study also reported that increased DNA methylation of POLG1 promoter could contribute to the metabolic memory phenomenon observed in the progression of diabetic retinopathy[42]. However, these studies are limited in that they haven’t detailed investigation in how metabolic memory alerts DNA methylation and the associated genes expression.

In this study, we found that elevated DNMT1 expression was involved in the sustained dysfunction of endothelial cells caused by metabolic memory. As one of DNA methyltransferases, in contrast to DNMT3a/3b, which mediate the establishment of new or de novo DNA methylation patterns [43], DNMT1 classically functions as a maintenance DNMT because it conserves the methylation pattern during replication [44]. Epigenetic modification by DNMT1 leads to various biological consequences related to inflammation, matrix remodeling and immune regulation [45]. Recently, several lines of evidence have shown that alterations in DNMT1 expression and activity play important roles in the diabetic complications [20], such as retinopathy [46], nephropathy [47] and cardiovascular complications [48]. Here we found that the expression and activity of DNMT1 were elevated in endothelial cells after transient hyperglycemia exposure, and inhibition of DNMT1 reversed the transient hyperglycemia-induced impaired angiogenesis, promoting wound healing, indicating that the increased DNMT1 is the key factor in the refractory wounds caused by metabolic memory. In addition, we also found that DNMT1 induces methylation in the Ang-1 promoter, resulting in decreased Ang-1 expression. As a strong endothelial-specific protective factor, Ang-1 promotes the formation of mature and functional microvessels, maintains endothelial integrity and protects microvessels against plasma leakage [49, 50]. Our previous study demonstrated that Ang-1 protects endothelial cells against advanced glycation end products (AGEs) damage, indicating the potential protective function of Ang-1 in microvascular injury in diabetic individuals [27]. Therefore, DNMT1-mediated Ang-1 hypermethylation served as a key mechanism in the impaired angiogenesis and delayed wound healing caused by metabolic memory. To further explore the underlying mechanisms by which DNMT1/Ang-1 signaling influences wound healing, we detected the activation of NF-κB. Previous studies have demonstrated that Ang-1 protects endothelial cells against hyperglycemic damage by inhibiting activation of the NF-κB pathway [33, 34], and the increased methylation and NF-κB activation also play important roles in diabetic complications [35, 36], suggesting that the NF-κB pathway may be involved in the impaired angiogenesis mediated by DNMT1/Ang-1 signaling. As expected, we found that activation of NF-κB was increased after transient hyperglycemia, and inhibition of DNMT1 or exogenous application of Ang-1 reversed transient hyperglycemia-induced NF-κB activation, indicating that DNMT1/Ang-1 affects angiogenesis and wound healing process by regulating the NF-κB pathway.

In summary, the observations reported here show that transient hyperglycemia causes persistent endothelial dysfunction during subsequent normoglycemia by inducing long-lasting changes in DNA methylation and recruitment of DNMT1 to the Ang-1 promoter, leading to decreased expression of Ang-1 and increased NF-κB activation. Together, these results implicate metabolic memory as an important mechanism in delayed wound healing in diabetes and indicate that blocking DNMT1 may have therapeutic value in treating diabetic vascular complications and delayed wound healing.

## Materials and Methods

### Cell culture

HUVECs were purchased from Lonza (Walkersville, MD) and maintained in DMEM (Sigma, St. Louis, MO) supplemented with 10% FBS and antibiotics. Cells at passages 3–5 were used and incubated in medium containing 30 mM high glucose (HG) or 5 mM normal glucose (NG) (Fig), and 30 mM mannitol served as an osmotic control.

### Animal experiments

#### Animals

Adult male C57B/6 mice were provided by the Experimental Animal Research Laboratory at Sun Yat-sen University in China. The animals were individually caged under specific pathogen-free (SPF) conditions with 12 h light-dark cycles. The animals were allowed access to water and standard rat chow ad libitum and were monitored daily. Treatment of the animals was carried out in strict accordance with the recommendations described in the Guide for the Care and Use of Laboratory Animals of the National Institutes of Health. The protocol was approved by the Committee on the Ethics of Animal Experiments of Sun Yat-sen University. All operations were performed under anesthesia, and all efforts were made to minimize suffering.

### Diabetic mouse protocol

The mice were divided into three groups: control group, diabetic group and metabolic memory group. Diabetes was induced in the mice with streptozotocin (55 mg/kg). In the diabetic group, the mice were maintained with poor glycemic control (glycated hemoglobin [GHb] was 10-12%) by receiving 1 unit of insulin (Humulin) every other day for 4 months. In the metabolic memory group, the mice were in poor glycemic control for 2 months, followed by good glycemic control (receiving insulin twice daily for a total of 8 units; GHb was 5-7%) for 2 additional months. Age-matched normal mice without any treatment served as the control group. The weights of the mice were measured twice a week, the blood glucose was detected once a week (Glucometer Elite), and GHb was evaluated every 2 months (Helena Laboratories).

### Induction of cutaneous wounds

The hair on the dorsal surface of the mice was removed by an electric shaver and cleaned with an alcohol swab. Two 6-mm-thick wounds were made on each side of the dorsum of the mice using a biopsy punch. siDNMT1 or 5-Aza diluted in PBS (30 ng/ml) was applied to the wounds, and PBS was used as a control. All wounds were covered with sterile gauze and bandaged. The experimental solutions were applied to the wounds daily, and the wounds were observed and assessed.

### Data analysis

All statistical analyses were performed using SPSS software, version 17.0 (SPSS Inc., Chicago, IL). Data in this study are expressed as the mean ± standard deviation (SD). The number of independent replicates of every part of an experiment is represented using the letter “n” in the figure legend. The differences between experimental groups were compared using the unpaired Student’s t-test or one-way ANOVA followed by post hoc analysis with the Bonferroni test. A P value of less than 0.05 was considered statistically significant.

## Acknowledgments

We would like to thank the Experimental Animal Research Laboratory of Sun Yat-sen University for providing technical supports for animals experiments. We also thank the Medical Science Experimentation Center of Sun Yat-sen University for providing the experimental facilities. We would also like to acknowledge the members of Centre for Translational Medicine of the first affiliated hospital of Sun Yat-sen University for helpful suggestions and discussions.

## Author Contributions

Bin Shu and Shaohai Qi designed experiments. Jingling Zhao, Shuai Yang and Lei Chen performed the in vitro experimental work and the histological detection. Ronghua Yang and Yingbin Xu performed the animal experiments and analysis. Julin Xie, Jun Wu and Xusheng Liu performed the data analysis. Jingling Zhao and Shuai Yang wrote the manuscript.

## Data and materials availability

All data associated with this study are present in the paper or the Supplementary Materials.

## REFERENCES AND NOTES

1. Laing T, Hanson R, Chan F, Bouchier-Hayes D. The role of endothelial dysfunction in the pathogenesis of impaired diabetic wound healing: a novel therapeutic target? Med Hypotheses. 2007; 69: 1029–31.

2. Ruttermann M, Maier-Hasselmann A, Nink-Grebe B, Burckhardt M. Local treatment of chronic wounds: in patients with peripheral vascular disease, chronic venous insufficiency, and diabetes. Dtsch Arztebl Int. 2013; 110: 25–31.

3. Dumville JC, Soares MO, O’Meara S, Cullum N. Systematic review and mixed treatment comparison: dressings to heal diabetic foot ulcers. Diabetologia. 2012; 55: 1902–10.

4. Edmonds M. Body of knowledge around the diabetic foot and limb salvage. J Cardiovasc Surg (Torino). 2012; 53: 605–16.

5. Falanga V, Eaglstein WH, Bucalo B, Katz MH, Harris B, Carson P. Topical use of human recombinant epidermal growth factor (h‐EGF) in venous ulcers. J Dermatol Surg Oncol. 1992; 18: 604–6.

6. Dinh TL, Veves A. The efficacy of Apligraf in the treatment of diabetic foot ulcers. Plast Reconstr Surg. 2006; 117: 152S–7S; discussion 8S‐9S.

7. Mustoe TA, O’Shaughnessy K, Kloeters O. Chronic wound pathogenesis and current treatment strategies: a unifying hypothesis. Plast Reconstr Surg. 2006; 117: 35S–41S.

8. Dinh T, Tecilazich F, Kafanas A, Doupis J, Gnardellis C, Leal E, et al. Mechanisms involved in the development and healing of diabetic foot ulceration. Diabetes. 2012; 61: 2937–47.

9. Diabetes C, Complications Trial Research G, Nathan DM, Genuth S, Lachin J, Cleary P, et al. The effect of intensive treatment of diabetes on the development and progression of long‐term complications in insulin‐dependent diabetes mellitus. N Engl J Med. 1993; 329: 977–86.

10. Turner RC, Cull CA, Frighi V, Holman RR, Grp UPDS. Glycemic control with diet, sulfonylurea, metformin, or insulin in patients with type 2 diabetes mellitus ‐ Progressive requirement for multiple therapies (UKPDS 49). Jama‐J Am Med Assoc. 1999; 281: 2005–12.

11. Holman RR, Paul SK, Bethel MA, Matthews DR, Neil HA. 10‐year follow‐up of intensive glucose control in type 2 diabetes. N Engl J Med. 2008; 359: 1577–89.

12. Kowluru RA. Effect of reinstitution of good glycemic control on retinal oxidative stress and nitrative stress in diabetic rats. Diabetes. 2003; 52: 818–23.

13. Roy S, Sala R, Cagliero E, Lorenzi M. Overexpression of fibronectin induced by diabetes or high glucose: phenomenon with a memory. Proc Natl Acad Sci U S A. 1990; 87: 404–8.

14. Li SL, Reddy MA, Cai Q, Meng L, Yuan H, Lanting L, et al. Enhanced proatherogenic responses in macrophages and vascular smooth muscle cells derived from diabetic db/db mice. Diabetes. 2006; 55: 2611–9.

15. Lee C, An D, Park J. Hyperglycemic memory in metabolism and cancer. Horm Mol Biol Clin Investig. 2016; 26: 77–85.

16. Brownlee M. The pathobiology of diabetic complications: a unifying mechanism. Diabetes. 2005; 54: 1615–25.

17. Giacco F, Brownlee M. Oxidative stress and diabetic complications. Circulation research. 2010; 107: 1058–70.

18. Pirola L, Balcerczyk A, Okabe J, El-Osta A. Epigenetic phenomena linked to diabetic complications. Nat Rev Endocrinol. 2010; 6: 665–75.

19. Fetita LS, Sobngwi E, Serradas P, Calvo F, Gautier JF. Consequences of fetal exposure to maternal diabetes in offspring. J Clin Endocrinol Metab. 2006; 91: 3718–24.

20. Reddy MA, Natarajan R. Epigenetic mechanisms in diabetic vascular complications. Cardiovasc Res. 2011; 90: 421–9.

21. Rodriguez H, El-Osta A. Epigenetic Contribution to the Development and Progression of Vascular Diabetic Complications. Antioxid Redox Signal. 2018; 29: 1074–91.

22. Reddy MA, Natarajan R. Role of epigenetic mechanisms in the vascular complications of diabetes. Subcell Biochem. 2013; 61: 435–54.

23. Maunakea AK, Chepelev I, Zhao K. Epigenome mapping in normal and disease States. Circulation research. 2010; 107: 327–39.

24. Kaufman PD, Rando OJ. Chromatin as a potential carrier of heritable information. Curr Opin Cell Biol. 2010; 22: 284–90.

25. Angst BD, Marcozzi C, Magee AI. The cadherin superfamily: diversity in form and function. Journal of cell science. 2001; 114: 629–41.

26. Vestweber D. Adhesion and signaling molecules controlling the transmigration of leukocytes through endothelium. Immunol Rev. 2007; 218: 178–96.

27. Zhao J, Chen L, Shu B, Tang J, Zhang L, Xie J, et al. Angiopoietin‐1 protects the endothelial cells against advanced glycation end product injury by strengthening cell junctions and inhibiting cell apoptosis. Journal of cellular physiology. 2015; 230: 1895–905.

28. Goldin A, Beckman JA, Schmidt AM, Creager MA. Advanced glycation end products: sparking the development of diabetic vascular injury. Circulation. 2006; 114: 597–605.

29. Cho CH, Kim KE, Byun J, Jang HS, Kim DK, Baluk P, et al. Long‐term and sustained COMP‐Ang1 induces long‐lasting vascular enlargement and enhanced blood flow. Circulation research. 2005; 97: 86–94.

30. Detich N, Hamm S, Just G, Knox JD, Szyf M. The methyl donor S‐Adenosylmethionine inhibits active demethylation of DNA: a candidate novel mechanism for the pharmacological effects of S‐Adenosylmethionine. The Journal of biological chemistry. 2003; 278: 20812–20.

31. Lu J, Song G, Tang Q, Zou C, Han F, Zhao Z, et al. IRX1 hypomethylation promotes osteosarcoma metastasis via induction of CXCL14/NF‐kappaB signaling. The Journal of clinical investigation. 2015; 125: 1839–56.

32. Lyko F. The DNA methyltransferase family: a versatile toolkit for epigenetic regulation. Nat Rev Genet. 2018; 19: 81–92.

33. He DK, Shao YR, Zhang L, Shen J, Zhong ZY, Wang J, et al. Adenovirus‐delivered angiopoietin‐1 suppresses NF‐kappaB and p38 MAPK and attenuates inflammatory responses in phosgene‐induced acute lung injury. Inhal Toxicol. 2014; 26: 185–92.

34. Fan F, Stoeltzing O, Liu W, McCarty MF, Jung YD, Reinmuth N, et al. Interleukin‐1beta regulates angiopoietin‐1 expression in human endothelial cells. Cancer Res. 2004; 64: 3186–90.

35. Zhang L, Zhang Q, Liu S, Chen Y, Li R, Lin T, et al. DNA methyltransferase 1 may be a therapy target for attenuating diabetic nephropathy and podocyte injury. Kidney Int. 2017; 92: 140–53.

36. Duraisamy AJ, Mishra M, Kowluru A, Kowluru RA. Epigenetics and Regulation of Oxidative Stress in Diabetic Retinopathy. Invest Ophthalmol Vis Sci. 2018; 59: 4831–40.

37. Pierce JW, Schoenleber R, Jesmok G, Best J, Moore SA, Collins T, et al. Novel inhibitors of cytokine‐induced IkappaBalpha phosphorylation and endothelial cell adhesion molecule expression show anti‐inflammatory effects in vivo. The Journal of biological chemistry. 1997; 272: 21096–103.

38. Park LK, Maione AG, Smith A, Gerami-Naini B, Iyer LK, Mooney DJ, et al. Genome‐wide DNA methylation analysis identifies a metabolic memory profile in patient‐derived diabetic foot ulcer fibroblasts. Epigenetics. 2014; 9: 1339–49.

39. Miao F, Smith DD, Zhang L, Min A, Feng W, Natarajan R. Lymphocytes from patients with type 1 diabetes display a distinct profile of chromatin histone H3 lysine 9 dimethylation: an epigenetic study in diabetes. Diabetes. 2008; 57: 3189–98.

40. Li Y, Reddy MA, Miao F, Shanmugam N, Yee JK, Hawkins D, et al. Role of the histone H3 lysine 4 methyltransferase, SET7/9, in the regulation of NF‐kappaB‐dependent inflammatory genes. Relevance to diabetes and inflammation. The Journal of biological chemistry. 2008; 283: 26771–81.

41. Olsen AS, Sarras MP, Jr., Leontovich A, Intine RV. Heritable transmission of diabetic metabolic memory in zebrafish correlates with DNA hypomethylation and aberrant gene expression. Diabetes. 2012; 61: 485–91.

42. Tewari S, Zhong Q, Santos JM, Kowluru RA. Mitochondria DNA replication and DNA methylation in the metabolic memory associated with continued progression of diabetic retinopathy. Invest Ophthalmol Vis Sci. 2012; 53: 4881–8.

43. Okano M, Bell DW, Haber DA, Li E. DNA methyltransferases Dnmt3a and Dnmt3b are essential for de novo methylation and mammalian development. Cell. 1999; 99: 247–57.

44. Robert MF, Morin S, Beaulieu N, Gauthier F, Chute IC, Barsalou A, et al. DNMT1 is required to maintain CpG methylation and aberrant gene silencing in human cancer cells. Nat Genet. 2003; 33: 61–5.

45. Robertson KD, Wolffe AP. DNA methylation in health and disease. Nat Rev Genet. 2000; 1: 11–9.

46. Kowluru RA, Shan Y, Mishra M. Dynamic DNA methylation of matrix metalloproteinase‐9 in the development of diabetic retinopathy. Lab Invest. 2016; 96: 1040–9.

47. Lu Z, Liu N, Wang F. Epigenetic Regulations in Diabetic Nephropathy. J Diabetes Res. 2017; 2017: 7805058.

48. Costantino S, Ambrosini S, Paneni F. The epigenetic landscape in the cardiovascular complications of diabetes. J Endocrinol Invest. 2018.

49. Yancopoulos GD, Davis S, Gale NW, Rudge JS, Wiegand SJ, Holash J. Vascular‐specific growth factors and blood vessel formation. Nature. 2000; 407: 242–8.

50. Lee SW, Kim WJ, Choi YK, Song HS, Son MJ, Gelman IH, et al. SSeCKS regulates angiogenesis and tight junction formation in blood‐brain barrier. Nat Med. 2003; 9: 900–6.

